# Protein Secondary Structure Patterns In Short-Range Cross-Link Atlas

**DOI:** 10.1101/2025.04.03.647081

**Authors:** Alice Vetrano, Alessio Di Ianni, Nico Di Fonte, Gianluca Dell’Orletta, Samantha Reale, Isabella Daidone, Claudio Iacobucci

**Author notes:** Human Technopole, Milan, Italy, V.le Rita Levi Montalcini 1, 20157, Milan (Alessio Di Ianni). These authors equally contributed (Alice Vetrano, Alessio Di Ianni).

## Abstract

Cross-linking mass spectrometry (XL-MS) has become a powerful tool in structural biology for investigating protein structure, dynamics, and interactomics. However, short-range cross-links, defined as those connecting residues fewer than 20 positions apart, have traditionally been considered less informative and largely overlooked, leaving significant data unexplored in a systematic manner. Here, we present a system-wide analysis of short-range cross-links, demonstrating their intrinsic correlation with protein secondary structure. We introduce the X-SPAN (Cross-link Structural Pattern Analyzer) software, which integrates publicly available XL-MS datasets from system-wide experiments with AlphaFold-predicted protein structures. Our analysis reveals distinct cross-linking patterns that reflect the spatial constraints imposed by secondary structural elements. Specifically, α-helices exhibit periodic cross-linking patterns consistent with their characteristic helical pitch, whereas coils and β-strands display nearly monotonic distributions. A context-dependent protein grammar reinforces short-range cross-link specificity. Short-range cross-links can enhance the statistical inference of secondary structures within integrative modeling workflows. Additionally, our work establishes a framework for benchmarking AlphaFold’s local prediction accuracy and provides novel quality control criteria for XL-MS experiments. We anticipate that X-SPAN and our short-range cross-link database will serve as a valuable resource for exploring local secondary structure rearrangements and their potential roles in protein function and allosteric regulation.

## Introduction

Structural biology aims to elucidate the three-dimensional structure of proteins and protein complexes as an essential step for understanding biological functions. Among the structural proteomics toolkit, cross-linking mass spectrometry (XL-MS) emerged as a key approach to study protein structures and interactions on a system-wide scale^[1–6]^. XL-MS comprehensively captures the morphology of dynamic biological assemblies in their native environment, providing structural data that enhances cryo-electron microscopy (cryo-EM) methods^[7, 8]^ and computational predictions. Given the relatively low structural resolution of XL-MS, typically a few tens of Å, the most informative insights come from cross-links between residues that are distant in the protein sequence or belong to different proteins. These cross-links provide valuable geometrical restraints, indicating the proximity of distinct regions, domains, and chains, thereby defining the global architecture of proteins. They help reveal long-range structural relationships and contribute to a deeper understanding of overall protein organization. Conversely, cross-links between residues that are close in sequence, e.g. fewer than 20 residues apart, are often considered less informative. Residues close in sequence are generally expected to be physically near each other in the three-dimensional (3D) space. Cross-links between such residues would impose trivial constraints that offer limited value in determining the 3D structure. The limited interest in short-range cross-links is reflected in how various XL-MS data analysis and visualization tools handle them. For instance, some XL search engines, such as MeroX^[9]^, offer the option to exclude all potential short cross-links from the analysis to reduce computational costs when performing proteome-wide studies. The important xiVIEW^[10]^-based PRIDE^[11]^ Crosslinking Database supports filtering cross-links based on a minimum residue separation to enhance visualization while preserving essential structural information.

Despite aligning with the common understanding that cross-links between distant residues in the sequence or between different proteins are fundamental in XL-MS, we hypothesize that short-range cross-links may still contain hidden structural information. Numerous XL-MS datasets from system-wide experiments are publicly available. In this study, we analyze short-range cross-links from these datasets by mapping them onto AlphaFold-predicted structures at a proteome-wide scale. Our analysis indicates that the spacing between identical residues depends on the local secondary structure and, to some extent, appears to be encoded at the genomic level. Upon normalization for this context-dependent amino acid distribution, clear secondary structure patterns emerge from experimental short-range cross-links. These patterns are determined by mutual orientation and spatial distances of side chains within distinct structural elements. Accurate interpretation of these observations requires precise geodesic measurements of solvent-accessible surface distances (SASD)^[12–17]^. Therefore, we refined existing computational tools to accurately represent the cross-linker pose. Moreover, the residue composition surrounding cross-linked sites reveals context-dependent characteristics, uncovering a specific protein grammar. We leveraged our short-range cross-link atlas to benchmark AlphaFold protein models across various predicted local distance difference test (pLDDT) scores, a per-residue measure of local confidence^[18, 19]^. Additionally, this atlas provides a novel quality-control framework for XL-MS experiments. Specifically, X-SPAN allows distinguishing genuine structural distributions from potential experimental artifacts.

## Results and Discussion

### Amino Acids Spacing in Secondary Structure Elements

The spacing between pairs of identical amino acids in secondary structural elements was examined across the proteomes of *Homo sapiens* (UP000005640), *Drosophila melanogaster* (UP000000803), and *Escherichia coli* (UP000000625) (Figures 1, S1-2). Secondary structural elements, specifically continuous α-helices, coils, β-strands, and mixed elements, were extracted from AlphaFold-predicted models of entire proteomes^[20]^ (Supporting Information). Subsequently, we computed the spacing between identical residues and analyzed their frequency distributions for all proteinogenic amino acids (Figures 1, S2). Our analysis revealed that genomes encode distinct preferred spacings for amino acid pairs, providing a glimpse into peptide folding upon translation. For instance, acidic and hydrophobic residues exhibit a characteristic spacing of approximately 3.6 residues within sequences encoding α-helices, reflecting helix amphipathicity. Additionally, distinct sequence motifs within secondary structural elements can be identified from residue pair distributions. Notably, the C2H2 zinc finger motif^[21]^ is characterized by a prominent HxxxH and CxxC peak in helices and coils, respectively (Figure 1). Another example is the ubiquitous K homology domain, marked by the conserved motif VIGxxGxxI^[22]^ within helices.

**Figure 1.**
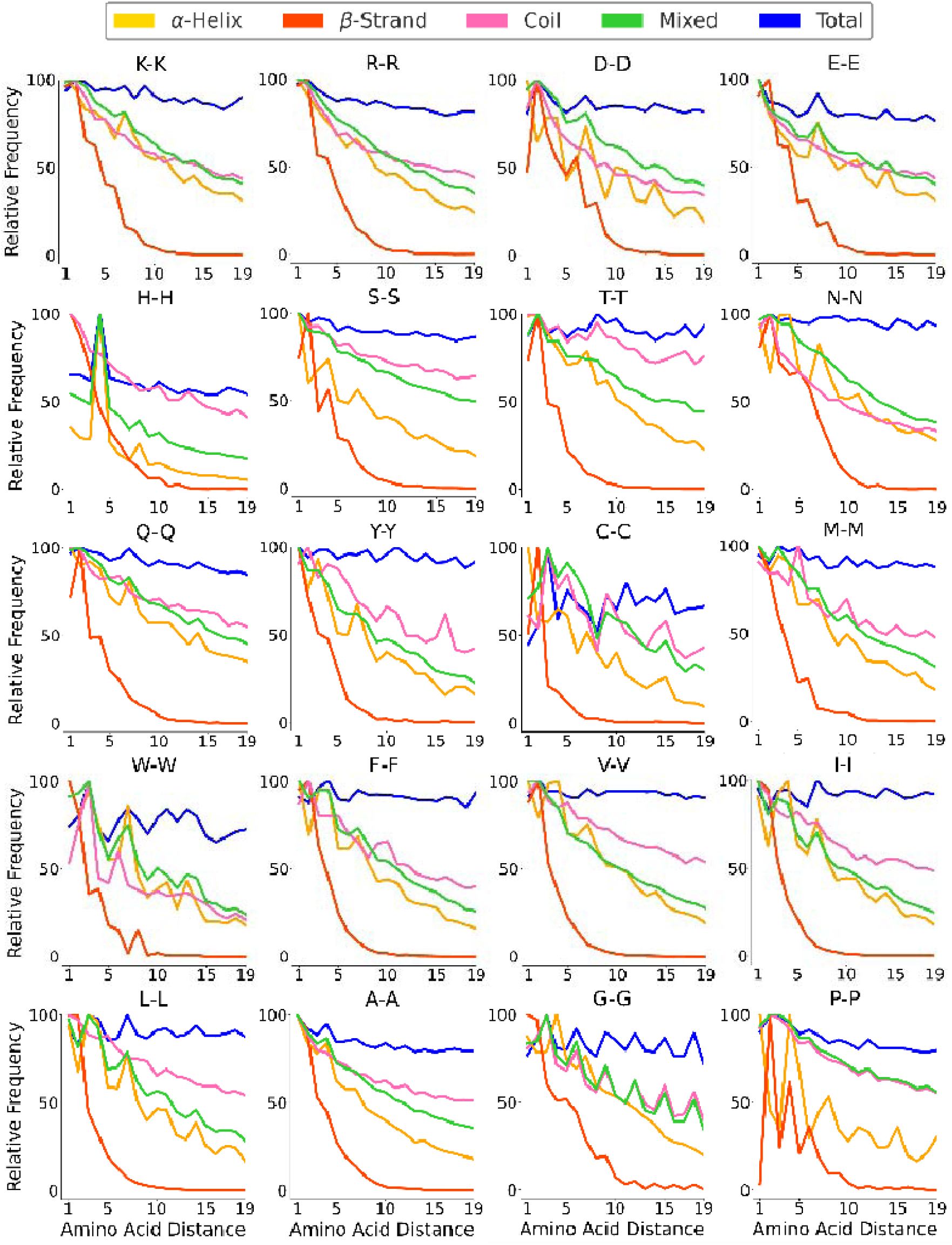
Relative frequency of pairwise amino acid distances across the human proteome. Each plot corresponds to an amino acid pair (e.g., K-K for lysine). The x-axis represents the amino acid spacing, while the y-axis represents the relative frequency normalized within each structural category. Curves are color-coded by secondary structure: α-helix (yellow), β-strand (red), coil (pink), mixed elements (green), and whole proteome (blue). Charged residues are plotted in the first row followed by polar and hydrophobic residues.

A central aspect of our analysis focuses on the spacing between lysine residues, the amino acids most frequently targeted in proteome-wide XL-MS experiments. Notably, lysine spacing does not display strongly distinct patterns across different secondary structural elements. Instead, it demonstrates a general decreasing frequency with increased residue spacing, aligning with the length distributions of continuous secondary structural elements within the proteome. This decline is particularly pronounced in β-strands, which typically consist of fewer residues due to their extended structure, and lysine residues are comparatively less abundant within β-strands compared to other elements (Figure S1). In contrast, α-helices, coils, and mixed elements share a more gradual frequency decline. We further investigated whether cross-linked lysine spacing deviates from this general proteomic trend. Such deviations could indicate differential cross-linker reactivity within secondary structural elements, rather than merely reflecting the underlying lysine distribution. Identifying these differences is essential for determining if cross-linking data offer additional structural insights beyond the general proteomic organization.

### Short-Range Cross-Link Analysis

We analyzed 12 publicly available system-wide XL-MS datasets^[9, 23–30]^ (Table S1), encompassing a total of 658,432 cross-links from four different organisms. These datasets include both amine-reactive and photo-reactive cross-linkers with varying spacer lengths. These 12 datasets were selected because each contained at least 400 unique short-range cross-links, a threshold that ensures sufficient sampling of the distance distribution. Cysteine-reactive cross-linkers were excluded from the analysis, as the presence of disulfide bonds cannot be reliably accounted for at the proteome-wide level, preventing a meaningful comparison between the experimental cross-links and the expected C–C distance distribution. Our analysis was carried out using X-SPAN, a Python-based software that annotates secondary structural elements for short-range cross-links based on AlphaFold protein models. X-SPAN accepts .mzIdentML 1.3, .zhrm, and .xlsx files as inputs and provides a graphical user interface (GUI) for ease of use. The workflow of the X-SPAN algorithm is outlined in Table 1, and the software is freely accessible at https://github.com/IacobucciLab/X-SPAN. X-SPAN categorizes all short-range cross-links into four groups (continuous α-helices, β-strands, and coils, and mixed secondary structure elements) and plots the distribution of cross-link frequencies as a function of reacted residue spacing. Results from amino-reactive and photo-activatable cross-linkers are presented in Figure 2 and Figure S3, respectively.

**Figure 2.**
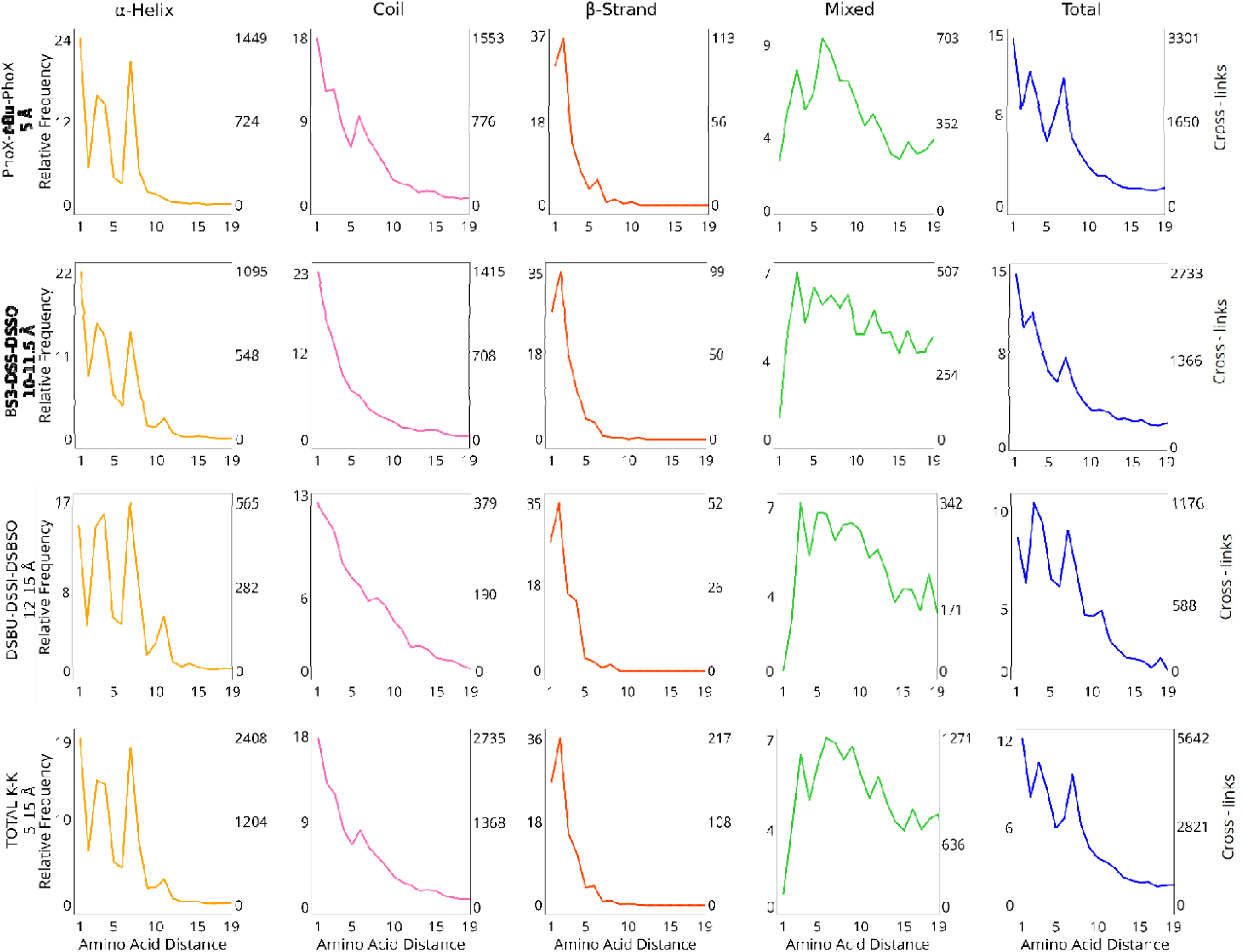
Relative frequency of K-K cross-links at increasing amino acid spacing. The x-axis represents the distance between amino acid pairs. The y-axis reports both the relative frequency of cross-links (left), normalized within each structural category to a maximum of 100, and the absolute counts (right). The four classes of structural elements are presented in separate columns and are color coded as follows: α-helix–yellow lines, β-strand–red lines, coil–pink lines, mixed elements–green lines, and total–blue lines. Distributions from cross-linkers with similar spacer length are presented in separate lines. Cumulative data are presented in the bottom row. Cross-link redundancy has been removed within each dataset.

**Table 1.**
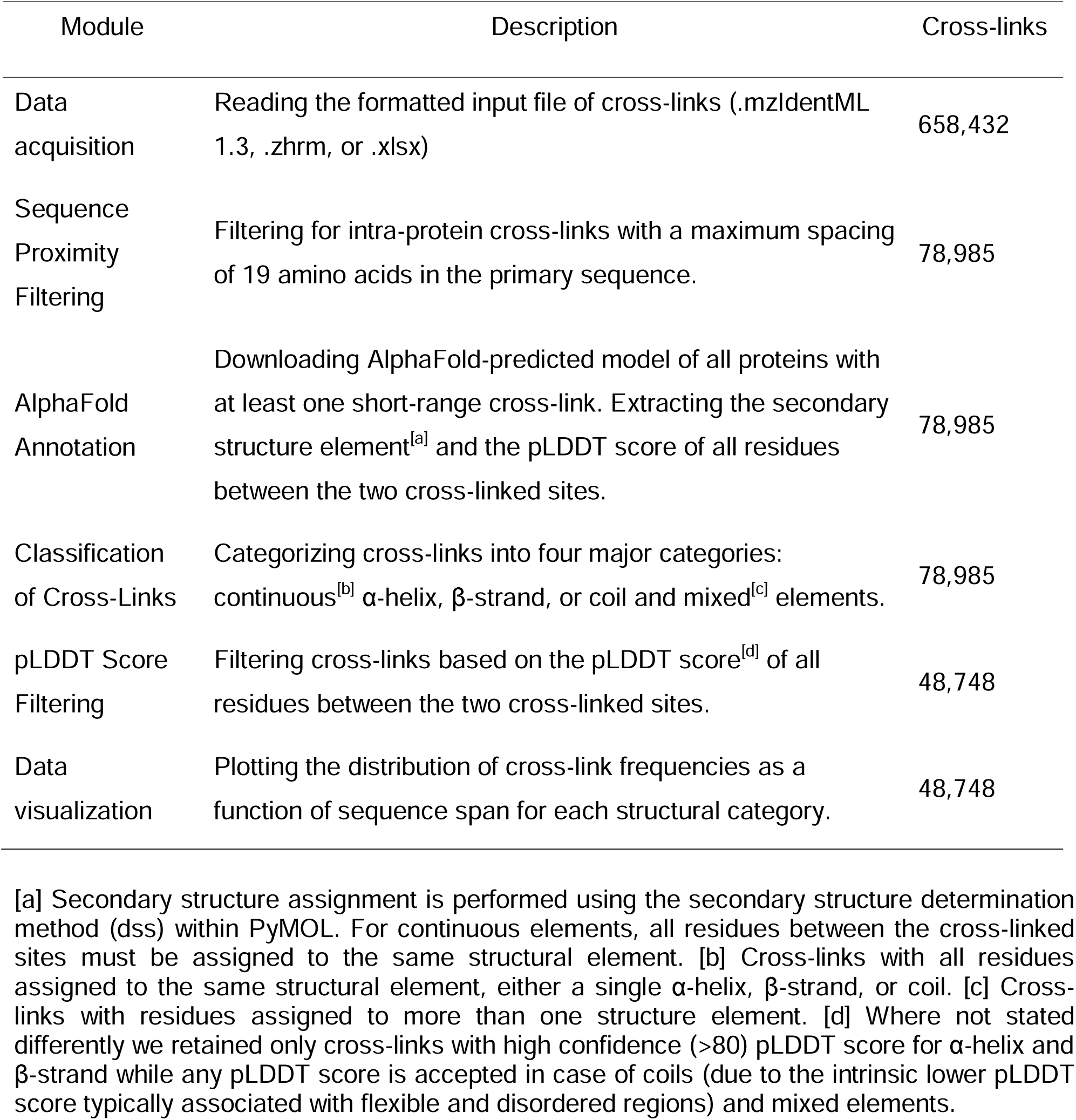
Concise description of X-SPAN workflow. Each module of the pipeline is described, along with the number of cross-links retained after each processing step.

#### α-helix

Among short-range cross-links, continuous α-helices represent the most common structural element (27%, Figure S1), independent of cross-linker reactivity or length. The distribution of cross-links by residue spacing strikingly differs from other secondary structures (Figures 2, S3) and from overall lysine spacing across the proteome (Figures 1, S2). Certain residue spacings are highly favored for cross-linking, while others appear almost prohibited. Their distribution follows an oscillating pattern modulated by a decreasing exponential trend (Figures 2, S3). The observed experimental periodicity of approximately 3.6 residues matches the helical pitch characteristic of α-helices. Meanwhile, the exponential curve’s slope reflects the length of the cross-linker utilized; shorter cross-linkers produce steeper declines due to their limited ability to reach distant residues. Conversely, longer cross-linkers can bridge residues across multiple helical turns without reducing spatial resolution. Surprisingly, shorter cross-linkers do not capture α-helix structures more effectively than longer cross-linkers, as the spacing between maxima and minima remains consistent across different linker lengths (Figures 2, S3). This may result from unfavorable entropic contributions associated with longer cross-linkers wrapping around helical structures ^[31]^.

We also examined patterns in non-specific photoactivatable cross-linkers^[26]^ (Figure S3). Such reagents provide high-density structural data^[32–34]^ independent of protein sequence, which could improve the detection of α-helix pattern compared to lysine-selective cross-linkers (Figure 2). Unexpectedly, although photocross-linkers still capture helical periodicity, they exhibit a reduction in the amplitude of the oscillations, evidenced by decreased distances between consecutive maxima and minima. This suggests that, for short-range cross-links, potential benefits of photoreactive cross-linkers might be counterbalanced by their lower precision in cross-linking site identification^[35, 36]^. While this lower accuracy has minimal implications for classical structural interpretations from XL-MS data, it becomes critical for identifying secondary structural elements through short-range cross-links.

#### Coils

Coils are protein secondary structure elements characterized by irregular and flexible shapes due to the absence of a consistent hydrogen-bonding network. Without a regular hydrogen-bonding pattern, the spacing between cross-linked residues lacks periodicity. Additionally, as the sequence span between cross-linked residues increases, the probability of cross-link formation declines (Figures 2, S3). Coils account for 33% of all short-range cross-links (Figure S1) and exhibit a gradual exponential decrease in cross-link frequency, reaching a maximum span of approximately 19 residues. Although coils are generally less compact than α-helices, their flexibility occasionally enables longer-range cross- links by bringing distant residues into proximity, albeit at much lower frequencies compared to more structured regions.

#### β-strand

Cross-links involving lysine residues in β-strands are notably less frequent compared to other structural elements. This is in agreement with previous reports in which low occurrence of lysine was reported for β-strands^[37, 38]^. Our findings show that short-range cross-links within continuous β-strands are even rarer, representing only 1% of the observed cases (Figure S1). The limited presence of lysine residues in β-strands (Figure S1) contributes significantly to their scarcity. Despite their low occurrence, specific preferred lysine distances are identifiable within experimental cross-links. In β-strands, side chains alternate orientation, projecting residues in opposite directions. This alternating pattern typically prohibits cross-link formation between residues with consecutive even-to-odd numbering. Therefore, a periodic distribution similar to that seen in α-helices could be expected, albeit with higher frequency. However, experimentally observed cross-links within β-strands exhibit a largely random distribution, decreasing rapidly with increased residue spacing, closely resembling the proteome-wide distribution (Figures 1, 2, S2, S3). This distribution pattern can be explained by the elongated and twisted nature of β-strands, where side-chain distances are non-uniform, enabling cross-linking at diverse spacings. Notably, a distinct peak occurs at a residue spacing of two, coinciding with optimal side-chain orientation despite the strand’s torsion. Interestingly, this spacing corresponds to a minimum observed in α-helix cross-link distributions.

#### Mixed elements

Cross-links within mixed structural elements exhibit a distinctive hump-shaped frequency distribution. Compared to other structural elements, their frequency is relatively low at short residue spacings but increases at intermediate distances before gradually decreasing. Unlike the distributions observed in other structural elements, cross-link frequency in mixed elements does not approach zero at longer spacings but stabilizes at a low plateau. At these extended distances, cross-links in mixed elements become predominant, likely due to compact structural motifs such as hairpins, loops, and turns, which bring distant residues into close spatial proximity. The short-range cross-link distribution (Figures 2, S3) differs significantly from overall lysine spacing within mixed elements at the proteome-wide level (Figures 1, S2). Notably, the occurrence of cross-links between consecutive residues strongly suggests these residues are part of either an α-helix or a coil.

Taken together, our results indicate that short-range cross-links serve as diagnostic markers for secondary structural elements. By analyzing the distribution of short-range cross-links obtained from all amino-reactive cross-linkers (Figure 2, bottom panel), we estimated the probabilities of peptides adopting specific secondary structures (Figure S4). For instance, at a spacing of two residues, the probability of the peptide forming a coil is 56%, nearly three times greater than the probabilities for α-helices or mixed elements. Conversely, at a residue spacing of seven, the likelihood of forming a continuous α-helix is approximately 50%, nearly double that observed for coils or mixed elements. Therefore, detecting short-range cross-links within an integrative modeling framework could significantly enhance inference of underlying peptide secondary structures.

To interpret these structural patterns (Figure 2), we refined the Biobox^[39]^ Python package to measure geodesic solvent-accessible surface distances (SASD) between reactive side-chain atoms (e.g., ε-nitrogen atom of lysine) while considering the cross-linker thickness (see Supporting Information). Commonly used Euclidean Cα-Cα distances systematically underestimate short-range cross-link distances due to neglecting surface curvature and local structural features (Figure S5, S6). Our SASD method more realistically represents the spatial pathways accessible to the cross-linker (Figures S5, S6). This geodesic approach effectively captures the periodic spacing characteristic of α-helices, aligning with experimental observations (Figures 2, S3, S5, S6).

### Context-Dependent Cross-Link Grammar

We also investigated whether the local amino acid environment affects the propensity of lysine to be cross-linked within different secondary structure elements. The composition of neighboring residues might promote or inhibit lysine cross-linking beyond structural constraints imposed by secondary structures. First, we analyzed the natural distribution of amino acids at varying distances around lysine within continuous secondary structures of the human proteome. We then compared these background distributions with those observed in our cross-link atlas (Figure 3). In α-helices, glutamic acid promotes lysine cross-linking when positioned at the first helical turn, at a spacing of three to four residues. At this distance, these residues reach the minimal geodesic separation optimal for hydrogen bonding, thus enhancing lysine nucleophilicity. This promoting effect is weaker for glutamine and aspartic acid due to their lack of charge and shorter side chain, respectively. Arginine, possessing a permanent positive charge, initially disfavors cross-linking within the first helical turn but reverses its effect beyond this region. Alanine facilitates lysine cross-linking, particularly when oriented towards the opposite face of the helix. Conversely, bulky hydrophobic residues, notably leucine, positioned within the first two helical turns, inhibit lysine cross-linking. Alanine and glutamic acid are also enriched around cross-linked lysine in coils, though without clear positional preferences. In coils, the inhibitory effects of certain residues, like leucine and cysteine, are weaker and less position-specific compared to α-helices. This aligns with the more extended and less structured nature of coils. Due to the limited experimental data available, the cross- link grammar of β-strands could not be fully characterized (Figure S7).

**Figure 3.**
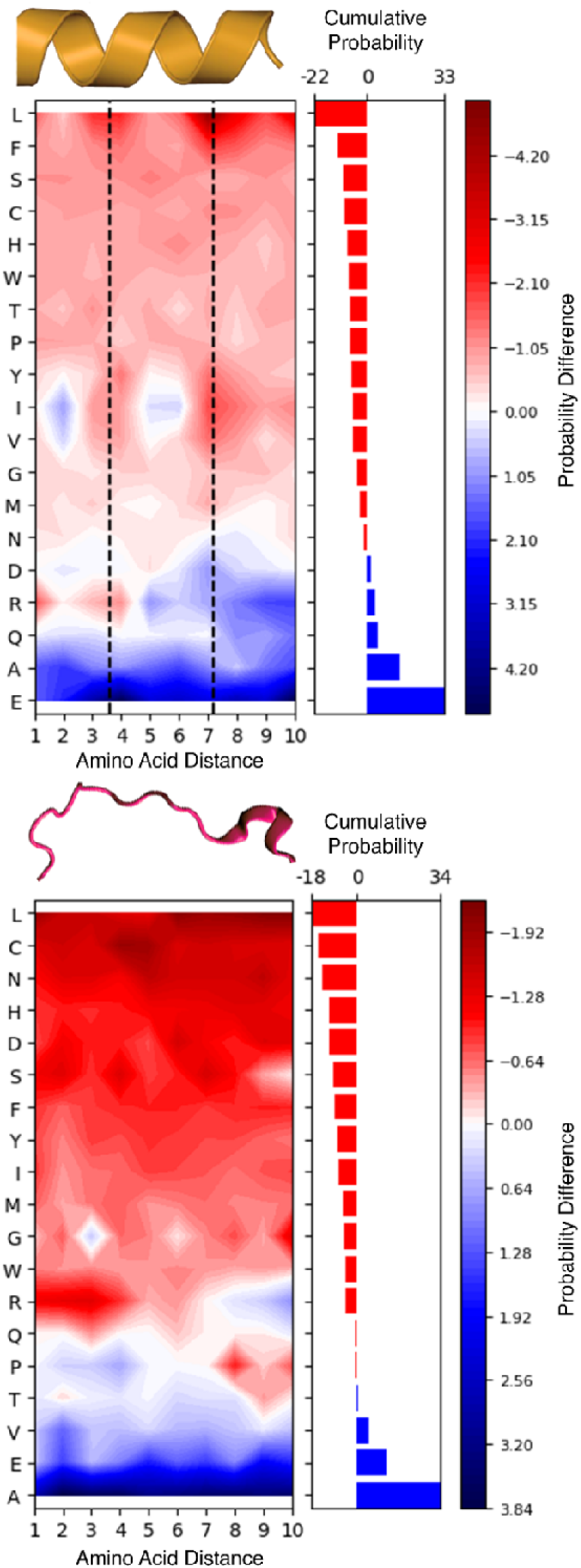
Influence of neighboring amino acids on lysine cross-linking propensity within α-helices (upper panel) and coils (bottom panel). Each heatmap represents the percentage difference in amino acid composition surrounding cross-linked lysine compared to the background proteome. The y-axis lists amino acids, while the x-axis indicates their sequence distance from a cross-linked lysine. Amino acids are ranked based on their overall effect on lysine cross-linking, with red indicating cross-linking inhibition and blue indicating cross-linking promotion. The right-side histogram illustrates this cumulative effect across all positions. Dashed lines mark the helical pitch, highlighting periodic cross-linking patterns in α-helices.

Finally, we analyzed the cross-link grammar distinguishing α-helices from coils (Figure 4). Residues positioned at the third, fourth, and seventh positions from lysine, corresponding to the first and second helical turns, emerged as diagnostic markers. Specifically, the presence of glutamic acid or aspartic acid at these positions indicates an α-helical structure, whereas hydrophobic residues typically suggest a coil structure.

**Figure 4.**
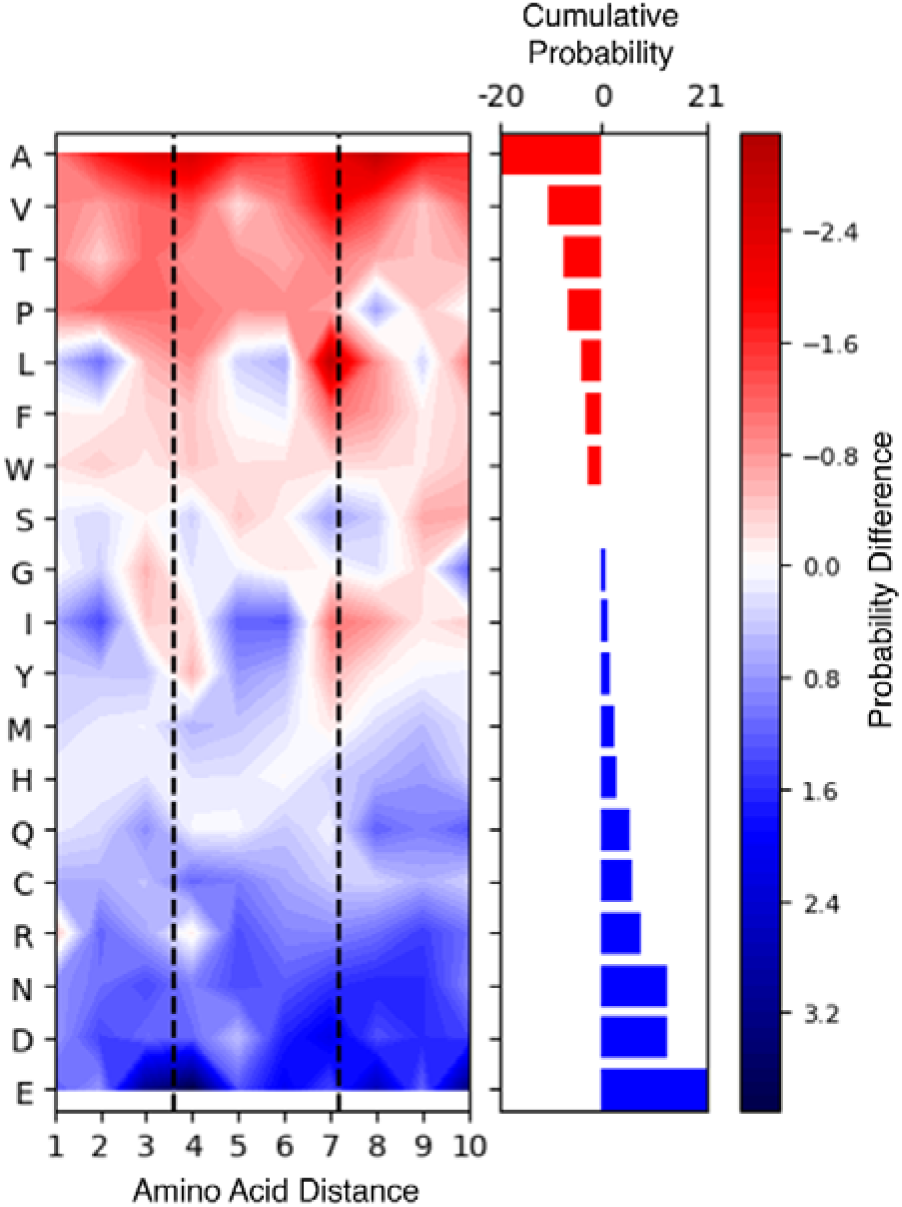
Influence of neighboring amino acids on cross-linked α-helix to coil recognition. The heatmap represents the percentage difference in amino acid composition between α-helices and coils. The y-axis lists amino acids, while the x-axis indicates their sequence distance from a cross-linked lysine. Amino acids are ranked based on their diagnostic capability, with red indicating residues more abundant in coils and blue indicating residues enriched in α-helices. The right-side histogram illustrates this cumulative effect across all positions. Dashed lines mark the helical pitch, highlighting periodic cross-linking patterns in α-helices.

The limited availability of experimental data prevented us from drawing conclusions from the comparison of the amino acid composition of β-strands versus coils, although this analysis is reported in Figure S8.

### Benchmarking AlphaFold-predicted secondary structure elements

Our short-range cross-link atlas provides a novel framework to evaluate the pLDDT score metric on a proteome-wide level. This investigation is based on structural data obtained in the native protein environment and is free from biases associated with the ability to express, purify, and determine high-resolution spectroscopic structures of proteins. We compare the cross-link distributions in secondary structure elements predicted within three pLDDT score ranges (0–60 for low local confidence, 60–80 for medium local confidence, and 80–100 for high local confidence) (Figure 5). The ranges are defined to ensure a sufficient number of cross-links for analysis. For the same reason, we combine all amine-reactive cross-linker datasets and exclude photo-reactive cross-linker datasets from this analysis.

**Figure 5.**
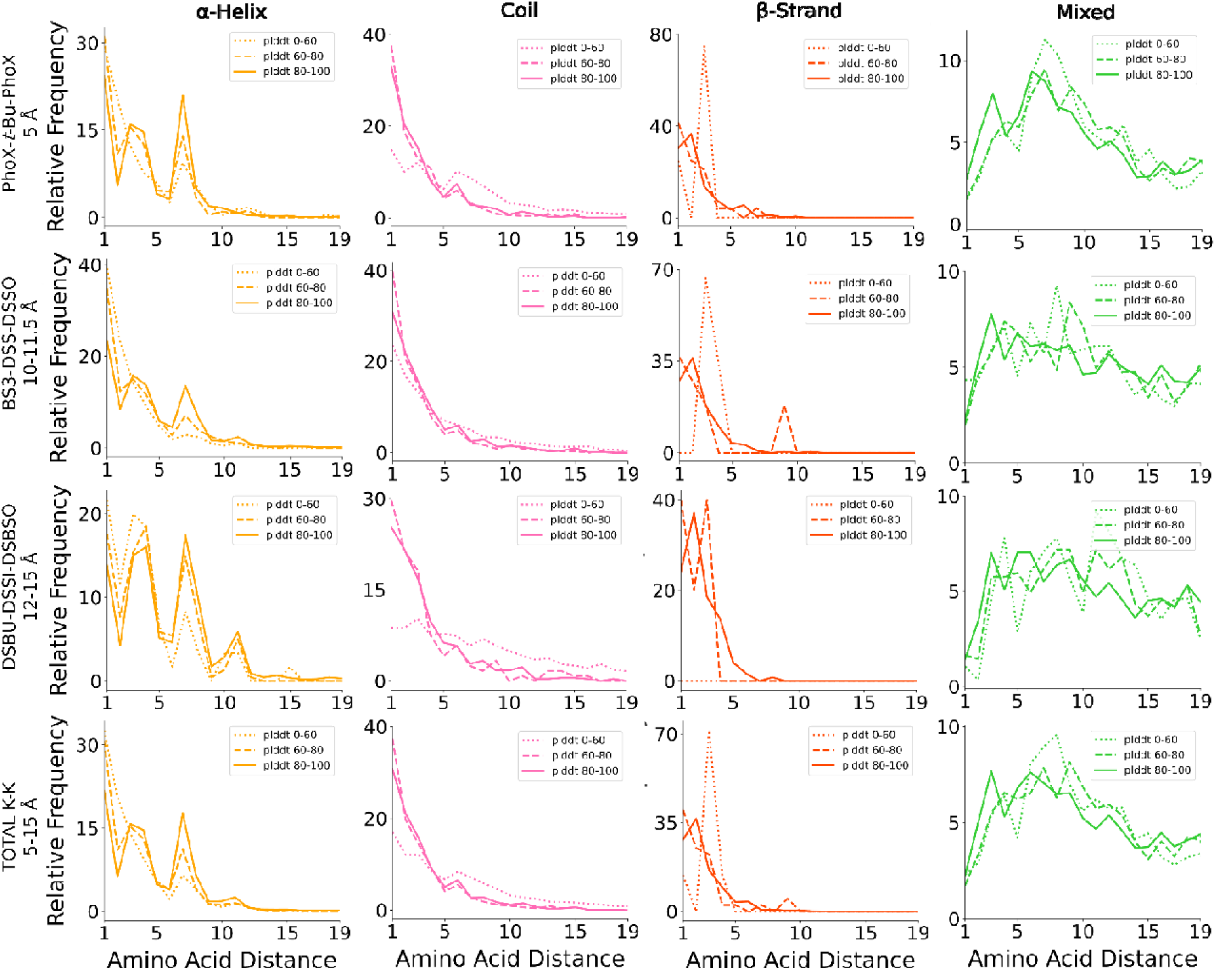
Distribution of K-K cross-links at different pLDDT scores. The x-axis represents the distance between amino acid pairs. The y-axis reports the relative frequency of cross-links, normalized within each structural category to a maximum of 100. The four classes of structural elements are presented in separate columns and are color coded as follows: α-helix–yellow lines, β-strand–red lines, coil–pink lines, and mixed elements–green lines. Distributions from cross-linkers with similar spacer length are presented in separate lines. Cumulative data are presented in the bottom row. pLDDT score ranges are presented as dotted line (0–60), dashed line (60–80), and solid line (80–100). Cross-link redundancy has been removed within each dataset.

Even at medium pLDDT confidence, the distinct patterns associated with α-helices were largely preserved. This indicates that α-helices with pLDDT scores greater than 60 are reliably predicted. Below this threshold, the α-helical pattern attenuated, and the cross-link distribution resembled that of coils with pLDDT scores above 60. This suggests that a significant portion of predicted α-helices at lower scores may be discontinuous or coil-like. For coils with pLDDT scores below 60, the cross-link distribution flattened, reflecting contributions from disordered regions with higher conformational flexibility, allowing cross-linking between more distant residues. β-strands appeared more sensitive to pLDDT scores, as their characteristic peak at a two-residue spacing disappeared at scores below 80, indicating that these segments might be less structured than predicted.

### Secondary Structure-Based Validation Metric

Validating cross-links in system-wide XL-MS experiments is critical to ensuring the reliability of structural insights. The most commonly used validation approach is structure-based validation, which evaluates cross-links by their agreement with known 3D structures of protein complexes. Although effective for targeted studies of individual proteins and complexes, this method has limitations for proteome-wide XL-MS experiments^[40]^. Alternatively, a STRING-based scoring approach has been proposed for validating protein-protein interaction (PPI) studies^[9]^. While meaningful, comparing experimentally detected PPIs against the background distribution in the STRING database does not provide a definitive validation criterion.

Here, we introduce a new metric based on the distribution of short-range cross-links within secondary structure elements as an additional layer of cross-link validation. We demonstrated that true positive short-range cross-links follow predictable patterns dictated by the spatial constraints of α-helices, β-strands, and coils (Figure 2, S3). In contrast, we expect false positive short-range cross-links to follow the natural occurrence of residue pairs within the proteome (Figure 1, S2). We hypothesized that datasets with a higher proportion of false positives, or those obtained under non-native conditions, would exhibit altered spacing distributions of short-range cross-links.

To test this hypothesis, we selected our recent DSSI^[28]^ cross-link dataset from HEK293T cell lysates (1% FDR at the PSM level) and generated three additional cross-link sets with drastically different quality levels. These sets were obtained by running the MeroX search engine using the original published settings but applying three progressively less stringent FDR criteria (1%, 5%, and 10%; see Supporting Information). We then applied X-SPAN, as defined in Table 1, obtaining 3681 short-range cross-links at 1% FDR, 17597 at 5% FDR, and 41422 at 10% FDR across the four categories of secondary structure elements (Figure 6). Notably, the distribution of lower-quality cross-link datasets, which contain a higher proportion of false positives, deviates significantly from that of the original 1% FDR dataset. Since false positives are randomly distributed, they cancel the characteristic α-helix periodic pattern and reduce the steepness of the20xponenttial decay observed in coils and β-strand. This behavior closely resembles what is observed at low-confidence pLDDT scores. These findings demonstrate that secondary structure-based validation of short-range cross-links is an effective approach for distinguishing datasets of varying quality. We propose that X-SPAN can provide an additional metric to define the overall quality of a proteome-wide cross-link dataset, which should align with the distribution patterns observed in our cross-link atlas.

**Figure 6.**
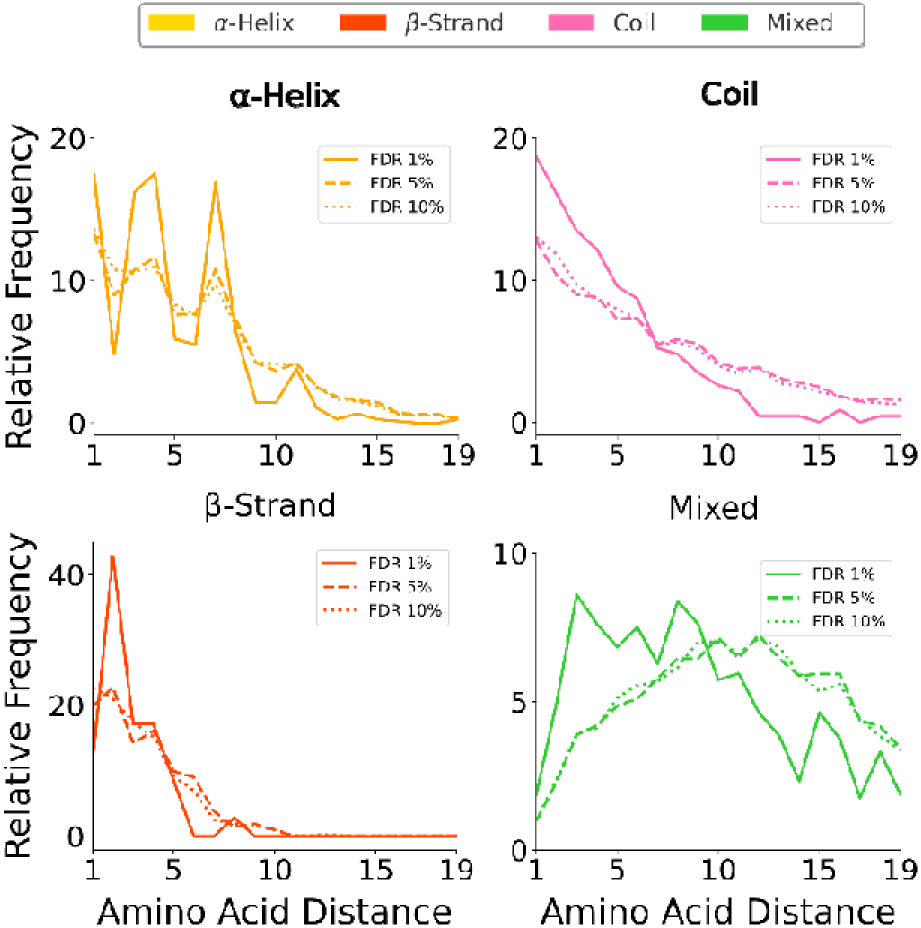
Distribution of DSSI cross-links at different FDR. The x-axis represents the distance between amino acid pairs. The y-axis reports the relative frequency of cross-links (left), normalized within each structural category to a maximum of 100. The four classes of structural elements are color coded as follows: α-helix–yellow lines, β-strand–red lines, coil–pink lines, and mixed elements–green lines. Different FDR thresholds are represented as dotted (10%), dashed (5%), and solid (1%) lines. Cross-link redundancy has been removed within each dataset.

## Conclusion

Short-range cross-links, traditionally considered less informative and frequently overlooked in XL-MS studies, have been shown here to carry intrinsic structural information. By systematically analyzing short-range cross-links across multiple proteomes, we demonstrated their strong correlation with secondary structural elements. Introducing X-SPAN, we integrated large, publicly available XL-MS datasets with AlphaFold-predicted proteomes, revealing distinct cross-linking patterns reflective of protein secondary structure organization. α-helices exhibit periodic cross-linking patterns matching their helical pitch, whereas coils and β-strands display nearly monotonic distributions influenced by their flexibility and extended conformation, respectively. Additionally, we identified a context-dependent protein grammar of cross-links, showing that local amino acid composition influences cross-linking probability. The prevalence of specific residues within secondary structures further supports the potential of short-range cross-links to assist secondary structure inference in integrative studies.

Our results demonstrate that short-range cross-links provide a useful benchmarking tool for evaluating AlphaFold’s local accuracy. By comparing experimental cross-link distributions with predicted structural elements across different pLDDT scores, we propose a data-driven method to assess AlphaFold model reliability at a local level. Furthermore, short-range cross- links introduce a novel quality control strategy for XL-MS experiments, distinguishing native structural distributions from potential artifacts.

The short-range cross-link atlas developed here, along with the broader application of our approach, may become a valuable resource for investigating secondary structure transitions and their roles in protein function and allosteric regulation.

## Supporting Information

The 12 datasets analyzed in this study were downloaded from the PRIDE database, and detailed information is provided in Table S1. The short-range cross-links extracted from all datasets are available as a .txt file. X-SPAN, along with all scripts used in this work, is freely accessible at https://github.com/IacobucciLab/X-SPAN.

## Acknowledgements

C.I. acknowledges financial support by the Italian Ministry of University and Research (MUR) (PRIN 2022─Project 20225HNCZK), by the European Union - NextGenerationEU (PRIN 2022 PNRR─Project P20224WAME), and by the European Union - NextGenerationEU under the Italian Ministry of University and Research (MUR) National Innovation Ecosystem grant ECS00000041 - VITALITY - CUP E13C22001060006.

