## Supplementary figures, tables, and methods for "Protein Secondary Structure Patterns In Short-Range Cross-Link Atlas"

[b] A. Di Ianni

Institution Human Technopole, Milan, Italy

V.le Rita Levi Montalcini 1, 20157, Milan

‡These authors equally contributed

*

**Table of contents**

**Material and Methods**

Public datasets ........................................................................................................ S-3

Amino acid pair distribution analysis ........................................................................... S-4

Geodesic SASD Measurements...................................................................................... S-4

MeroX settings…............................................................................................................. S-5

**Additional Figures**

Figure S1 ........................................................................................................................ S-5

Figure S2 ........................................................................................................................ S-6

Figure S3 ........................................................................................................................ S-7

Figure S4 ........................................................................................................................ S-8

Figure S5 ........................................................................................................................ S-9

Figure S6 ........................................................................................................................ S-10

Figure S7 ........................................................................................................................ S-11

Figure S8 ........................................................................................................................ S-12

**References**

....................................................................................................................................... S-13

### Material and Methods

#### Public datasets

**Table S1.** Summary of cross-link datasets[1-9] analyzed, detailing the cross-linkers used (XL), the protein source organisms (Organism), the corresponding authors (Corr. Author), the publication year (Year), the software used for analysis (Software), and the total number of cross-link spectral matches (CSMs) identified. The data span from 2019 to 2024, featuring various cross-linking search software, including MeroX, XlinkX, xiSEARCH, pLink, and Protein Prospector.

| *Entry* | *XL* | *Organism* | *Corr. Author* | *Year* | *Software* | *CSMs* | *Unique <20AA* |
|---|---|---|---|---|---|---|---|
| ***1*** | DSBU[a] | Drosophila | A. Sinz | 2019 | MeroX 2.0 | 56,402 | 8,775 |
| ***2*** | DSSO | Mus musculus | A. B. Smit | 2020 | XlinkX 2.0 | 14,319 | 3,913 |
| ***3*** | BS3/ DSS | Ecoli | J. Rappsilber | 2021 | xiSEARCH 1.6.746 | 42,247 | 2,584 |
| ***4*** | *t*-Bu-PhoX | Human | F. Liu | 2022 | pLink 2.3.9 | 16,152 | 14,690 |
| ***5*** | dizSEC | Ecoli | S. D. Fried | 2023 | XiSEARCH 1.7.6.7 | 1,923 | 563 |
| ***6*** | dizSPC | Ecoli | S. D. Fried | 2023 | XiSEARCH 1.7.6.7 | 1,279 | 420 |
| ***7*** | DSBSO | Human | L. Huang | 2024 | Protein Prospector 6.3.5 | 13,227 | 1,017 |
| ***8*** | DSBU [a][b] | Human | C. Iacobucci | 2024 | MeroX 2.0 | 163,790 | 13,151 |
| ***9*** | DSSI [a] | Human | C. Iacobucci | 2024 | MeroX 2.0 | 42,781 | 3,330 |
| ***10*** | DSS | Human | D. C. Schriemer | 2024 | pLink 2.3.11 | 148,194 | 14,605 |
| ***11*** | PhoX | Human | D. C. Schriemer | 2024 | pLink 2.3.11 | 148,967 | 10,155 |
| ***12*** | *t*-Bu-PhoX | Human | J.S. Choudhary | 2024 | pLink 2.3.11 | 9,151 | 5,119 |

[a] These datasets originally obtained by MeroX have been reanalyzed using MeroX 2.0.1.7 software to include loop-links in the search. We downloaded and applied the original FASTA file and MeroX settings of each dataset as indicated in the original publication exept for enabling “intrapeptidal cross-links (type I)”.

[b] A score cut off of 80 or sum pepscore cutoff of 150 has been applied upon spectra inspection.

**Amino acid pair distribution analysis**

The Python script developed to analyze the distance distribution of amino acid pairs in secondary structure elements is freely availbale at https://github.com/IacobucciLab/X-SPAN

**Geodesic SASD Measurements**

To further rationalize the capability of cross-linkers to react differently within secondary structure elements, we measure the actual distance of residues in the 3D space. The majority of XL-MS studies applies the euclidean metric, typically considering the Cα atoms. This approach is widely accepted and provides a valid approximation of the distance that the cross-linker must cover when bridging different structural elements, protein regions, and different proteins. We find that the conventional Cα-Cα distance metric becomes inadequate for short-range cross-links where the reactive residues are close in sequence. The Cα-Cα distance leads to a systematic underestimation of the actual distance that the cross-linker must cover (Figure S5, S6). This is due to the heterogeneity of amino acid spacing of different secondary structure elements. An α-helix is more compact than a coil or a β-strand, meaning that two lysine positioned a few residues apart in sequence will exhibit a low Euclidean Cα-Cα distance that is not representative of the actual spacing between their reactive amine groups. In the literature, several tools are available to measure Euclidean or solvent-accessible surface distances (SASD) between Cα, Cβ, and ε-amino groups[10, 11]. While being more accurate, we found that also these tools tend to underestimate the true distance between two ε-amino groups in short-range cross-links. Even SASD-calculating tools that attempt to correct for solvent accessibility often approximate distances using Cα atoms, which do not lie directly on the solvent-accessible surface. Others compute SASD by approximating it as a one-dimensional geodesic path along the solvent-accessible surface. However, the cross-linker has a finite thickness that further increases the SASD. To estimate the SASD for short-range cross-links, we modified Biobox[12] Python package. In particular, we measure the shortest geodesic SASD while accounting for cross-linker thickness. To achieve this, we double the atomic radius of the atoms used to define the SASA in Biobox, effectively incorporating the cross-linker's atomic radius. This adjustment improves the approximation of the cross-linker pose (Figure S5, S6), aligning better with experimental cutoffs observed in α-helices (Figure 2, S3). To illustrate our approach, we apply it to model poly-lysine and poly-glycine α-helices (Figure S6), capturing differences in steric hindrance.

**MeroX settings**

The utility of X-SPAN as a quality control tool for XL-MS experiments was evaluated using the DSSI[7] dataset. This dataset was reanalyzed with MeroX, applying false discovery rate (FDR) cutoffs of 1%, 5%, and 10% at the PSM level, while all other settings were kept consistent with the original publication[7].

**Additional Figures**

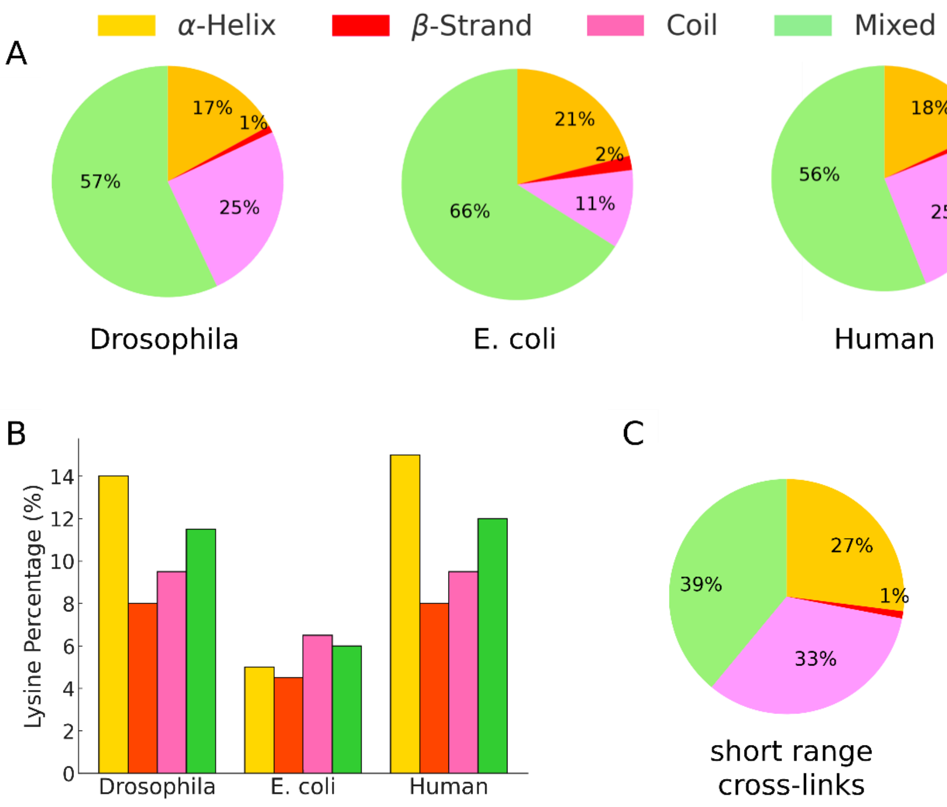


**Figure S1.** (A) Pie charts showing the global distribution of lysine across different secondary structure elements in various proteomes. (B) Bar chart depicting the percentage of lysines relative to other amino acids within different secondary structure elements and across organisms. (C) Pie chart illustrating the experimental distribution of short-range cross-linked lysine among secondary structure elements of the human proteome.

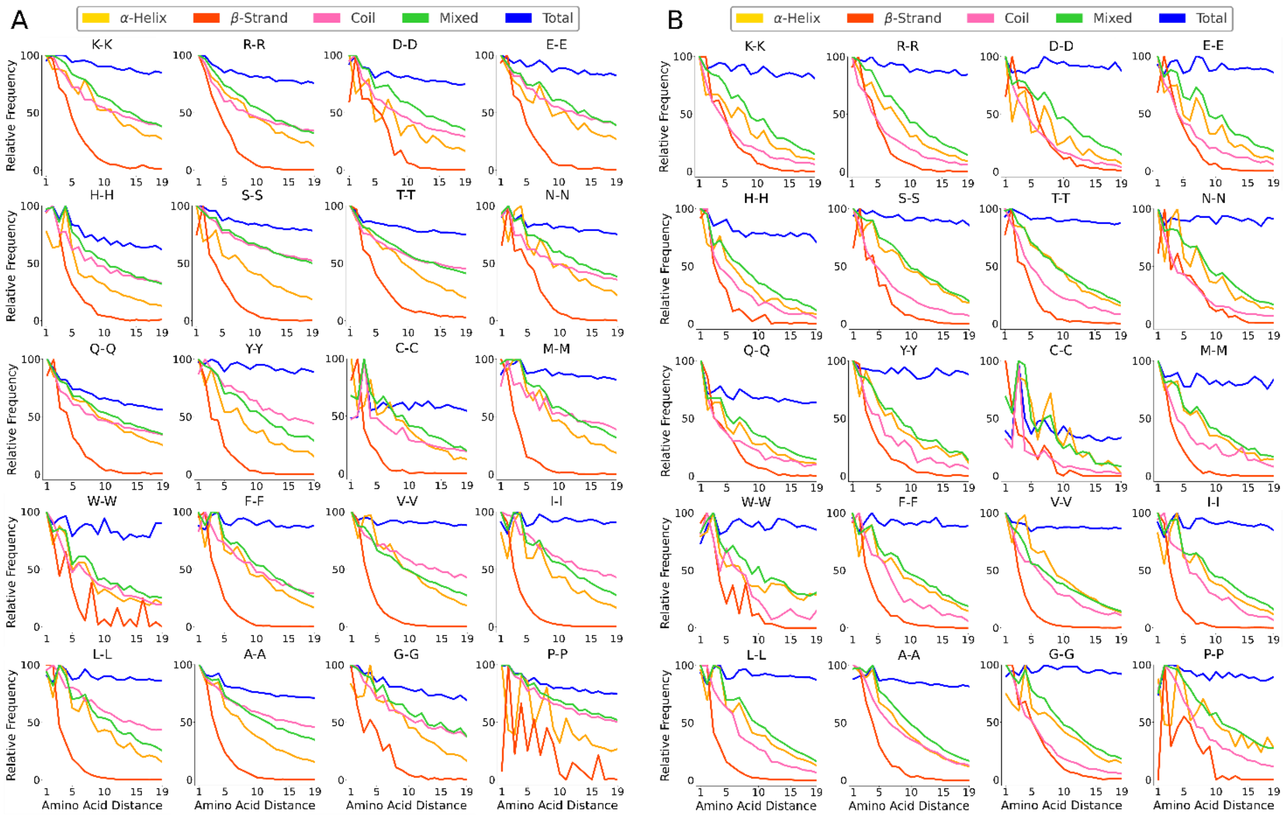


**Figure S2.** Relative frequency of pairwise amino acid distances across Drosophila (A) and E.coli (B) proteomes. α-helix (yellow lines), β-strand (red lines), coil (pink lines), mixed elements (green lines), and the entire proteome (blue lines). Each plot corresponds to an amino acid pair (e.g., K-K for lysine). The x-axis represents the amino acid spacing, while the y-axis represents the relative frequency normalized within each structural category. Charged residues are plotted in the first row followed by polar and hydrophobic residues.

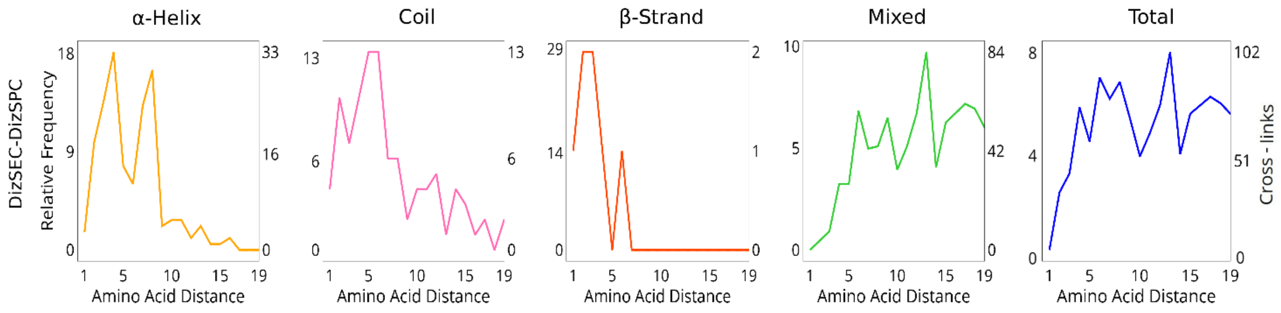


**Figure S3.** Relative frequency of short-range K-X cross-links at increasing amino acid spacing obtained with dizSEC and dizSPC photocross-linkers[5].The x-axis represents the distance between amino acid pairs. The y-axis reports both the relative frequency of cross-links, normalized within each structural category to a maximum of 100, and the absolute counts. The four classes of structural elements are presented in separate columns and are color coded as follows: α-helix–yellow lines, β-strand–red lines, coil–pink lines, mixed elements–green lines and total–blue lines. It is worth noting that the very low number of cross-links observed between consecutive residues (amino acid distance = 1) is due to the absence of loop-links in the deposited list of identified cross-links. As a result, cross-links between adjacent residues are underrepresented. When the first of these two cross-linked residues is a lysine, it is unlikely that trypsin cleaves the peptide bond between them, since the lysine side chain loses its positive charge upon cross-linking, reducing recognition by the protease.

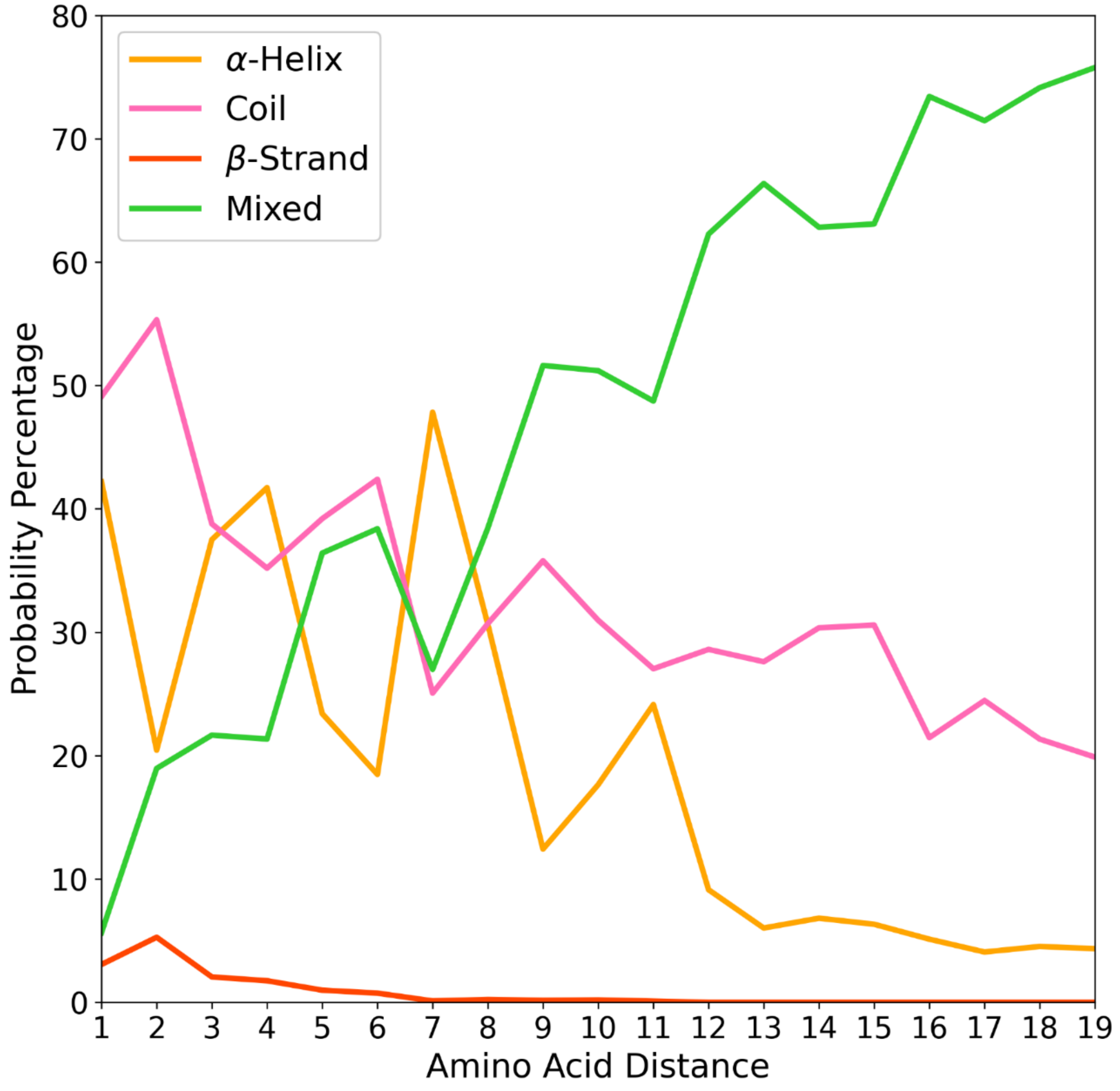


**Figure S4.** Probability distribution of secondary structure elements at different cross-link distances. The probabilities have been calculated considering all short-range cross-links obtained from amino-reactive cross-linkers (Figure 2, bottom panel). The stacked bar plot represents the probability percentage of different secondary structure elements (α-helices (yellow), coils (pink), β-strands (red), and mixed structures (green)) at varying cross-link sequence distances. The x-axis represents the sequence distance between cross-linked residues, while the y-axis indicates the probability percentage of each secondary structure type.

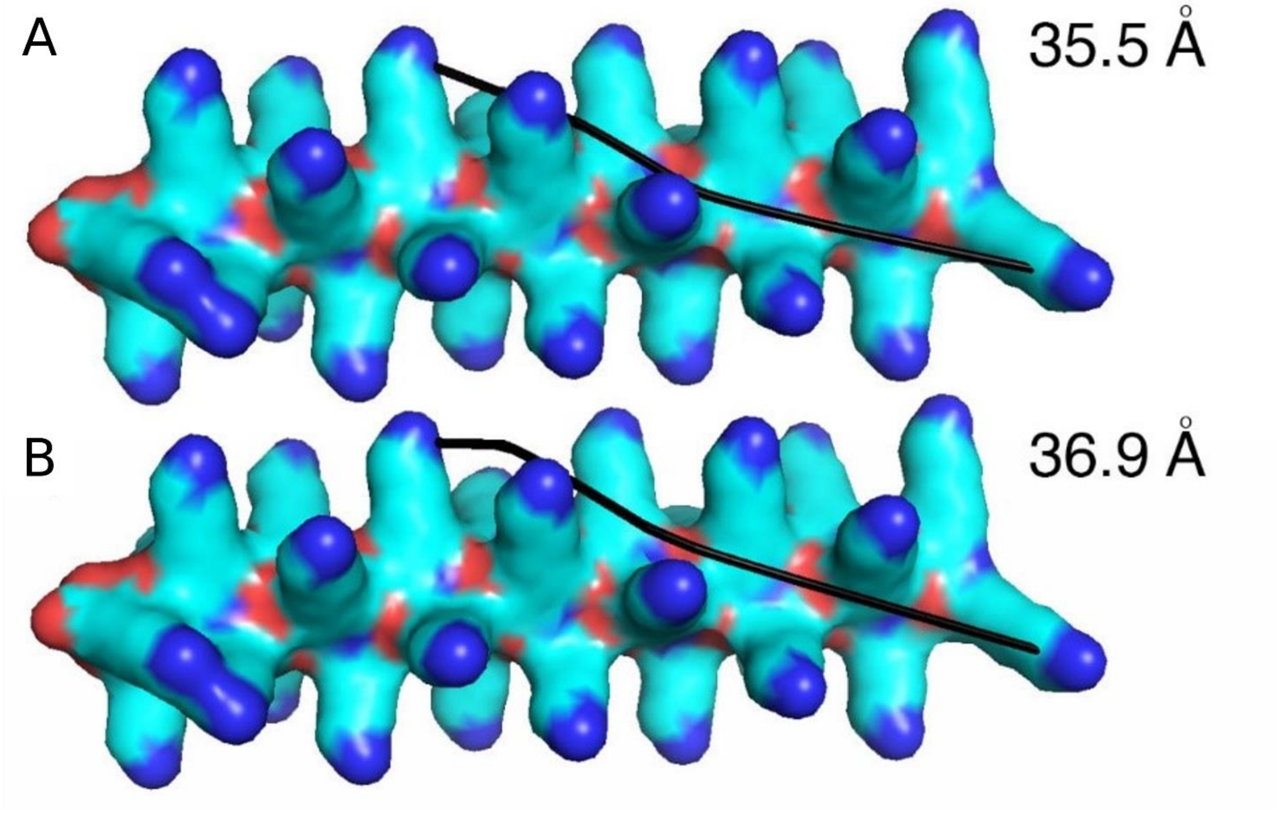


**Figure S5:** Comparison of geodesic distances measured with a) standard Biobox b) Biobox with a doubled clashing radius in a model poly-lysine α-helix. The reactive ε-nitrogen atoms of lysine were considered for measuring the distances. Measuring geodesic distances with Biobox determines a slight underestimation of amino group distance. Implicitly considering the thickness of the cross-linker, by adjusting the clash radius by doubling its value, improve the cross-link distance estimation (see also figure S6).

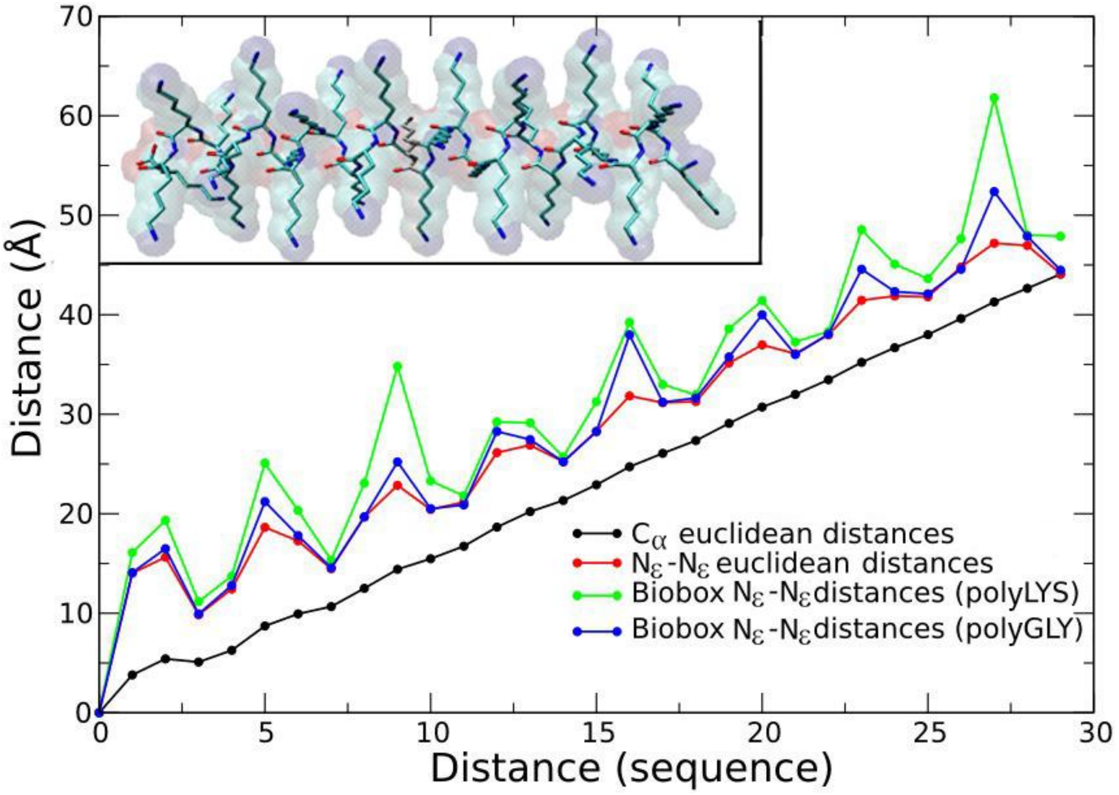


**Figure S6:** Comparison of Euclidean and geodesic cross-link distances measured with modified Biobox in a model poly-lysine and poly-glycine α-helix. The reactive ε-nitrogen atoms of lysine were considered for measuring the distances. In the poly-glycine peptide, two residues at each evaluated distance were replaced by lysine to enable the amino group distance measurement.

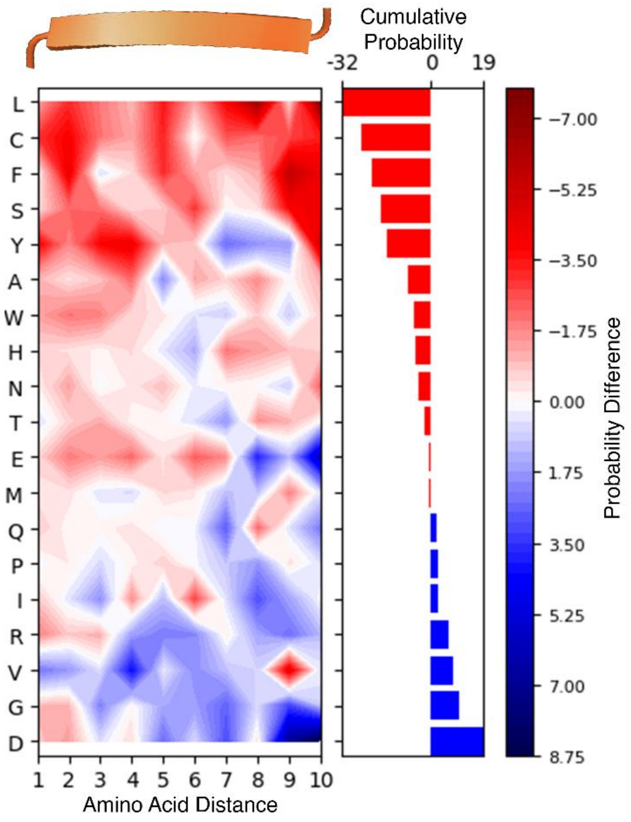


**Figure S7.** Influence of neighbouring amino acids on lysine cross-linking propensity within β-strands. The heatmap represents the percentage difference in amino acid composition surrounding cross-linked lysine compared to the background proteome. The x-axis represents the sequence distance between cross-linked residues. The y axis displays different amino acids, ordered by their cumulative probability of influencing lysine cross-linking. The right histogram illustrates the cumulative effect of each amino acid, summing its influence across the first ten residues around lysine. Bars in blue denote amino acids that globally enhance lysine cross-linking, whereas red bars indicate inhibitory effects. Colors indicate the probability of promoting (blue) or inhibiting (red) lysine cross-linking. The upper plot summarizes the average effect of each sequence position across all amino acids, providing an overall trend of how amino acid positioning influences lysine cross-linking propensity.

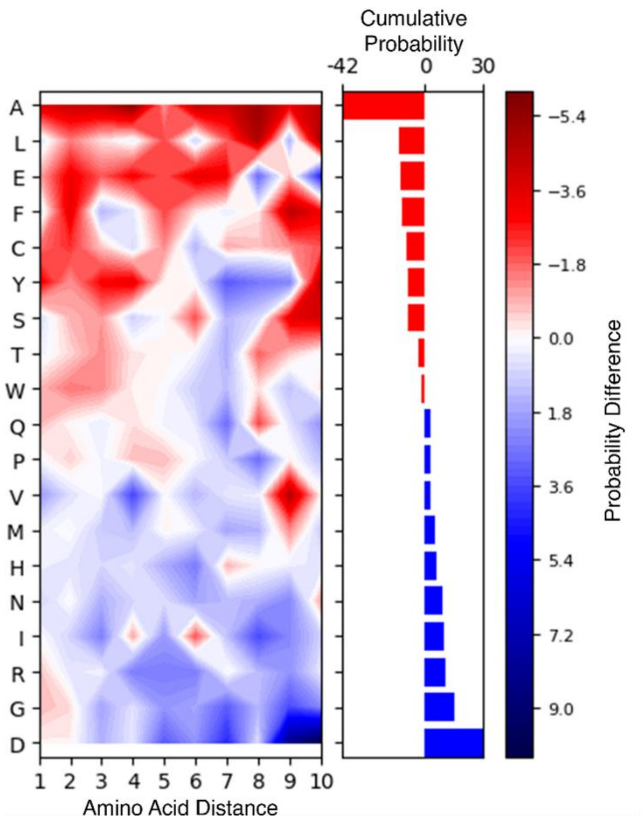


**Figure S8.** Influence of neighboring amino acids on cross-linked β-strand to coil recognition. The heatmap represents the percentage difference in amino acid composition between β-strands and coils. The x-axis represents the sequence distance between cross-linked residues. The y axis displays different amino acids, ordered by their cumulative diagnostic capability. The right histograms illustrate the cumulative effect of each amino acid, summing its differential abundance between β-strands and coils over the first ten residues around cross-linked lysine. Bars in blue denote amino acids that are globally enriched in β-strands, whereas red bars indicate residues more abundant in coils. Colors indicate the probability of a sequence adopting a β-strand fold (blue) or a coil fold (red). The upper plots summarize the average effect of each sequence position across all amino acids, providing an overall trend of how amino acid positioning influences the diagnostic power of the fold.
